# Dehydration risk, not ambient incubation, limits nest attendance at high temperatures

**DOI:** 10.1101/2020.11.18.388025

**Authors:** Amanda R. Bourne, Amanda R. Ridley, Andrew E. McKechnie, Claire N. Spottiswoode, Susan J. Cunningham

**Affiliations:** FitzPatrick Institute of African Ornithology, DST-NRF Centre of Excellence, University of Cape Town, Private Bag X3, Rondebosch 7701, South Africa; Centre for Evolutionary Biology, School of Biological Sciences, University of Western Australia, Crawley 6009, Australia; South African Research Chair in Conservation Physiology, South African National Biodiversity Institute, Pretoria, South Africa; DST-NRF Centre of Excellence at the FitzPatrick Institute, Department of Zoology and Entomology, University of Pretoria, Hatfield, South Africa; Department of Zoology, University of Cambridge, Downing Street, Cambridge CB2 3EJ, UK

**Keywords:** Climate change, cooperative breeding, high temperatures, incubation, parental care, southern pied babbler

## Abstract

High air temperatures have measurable negative impacts on reproduction in wild animal populations, including during incubation in birds. Understanding the mechanisms driving these impacts requires comprehensive knowledge of animal physiology and behaviour under natural conditions. We used a novel combination of a non-invasive doubly-labelled water technique and behaviour observations in the field to examine effects of temperature, rainfall, and group size on physiology and behaviour during incubation in southern pied babblers *Turdoides bicolor*, a cooperatively-breeding passerine endemic to a semi-arid region in southern Africa. The proportion of time that clutches of eggs were incubated declined as air temperatures increased, traditionally interpreted as a benefit of ambient incubation. However, we show that a) clutches were less likely to hatch when exposed to high air temperatures; b) pied babbler groups incubated their nests almost constantly (97% of daylight hours) except on hot days; c) operative temperatures in unattended nests were substantially higher than air temperatures and frequently exceeded 40.5°C, above which bird embryos are at risk of death; d) pied babblers incubating for long periods of time failed to maintain water balance on hot days but not cool days; and e) pied babblers from incubating groups did not maintain body mass on hot days. These results suggest that, rather than taking advantage of opportunities for ambient incubation, pied babblers leave the nests during hot periods to avoid dehydration as a consequence of incubating at high operative temperatures. As mean air temperatures increase and extreme heat events become more frequent under climate change, birds will likely incur greater water costs during incubation, leading to compromised nest attendance and increased likelihood of eggs overheating, with implications for nest success and, ultimately, population persistence.

## Introduction

Anthropogenic climate change is driving population declines in birds globally (Saino *et al.*, 2011; Iknayan and Beissinger, 2018; Rosenberg *et al.*, 2019), often associated with negative impacts on reproduction (Stevenson and Bryant, 2000; Cahill *et al.*, 2013; Cunningham *et al.*, 2013). For example, hatching failure in birds is particularly common during hot weather (Wada *et al.*, 2015; Clauser and McRae, 2017) and droughts (Conrey *et al.*, 2016), both of which are becoming more frequent under climate warming (Ripple *et al.*, 2019). Understanding the behavioural and physiological mechanisms driving such patterns *in situ* in wild populations is critical to our ability to predict species’ responses to climate change (Conradie *et al.*, 2019; Stillman, 2019).

Incubation is energetically costly in temperate environments where eggs need to be kept warm (Ardia *et al.*, 2010; Nord *et al.*, 2010; Nord and Cooper, 2020), but also extremely challenging in warm environments (Amat and Masero, 2004; Coe *et al.*, 2015; Nwaogu *et al.*, 2017), where incubating birds must prevent eggs from overheating (Grant, 1982; Carroll *et al.*, 2015; McDonald and Schwanz, 2018) while also thermoregulating themselves (DuRant *et al.*, 2019; McKechnie, 2019). Birds initially respond to high air temperatures (T_air_) by increasing incubation constancy (AlRashidi *et al.*, 2011; Mortensen and Reed, 2018) or engaging in shading behaviours (Grant, 1982; Downs and Ward, 1997; Brown and Downs, 2003; Clauser and McRae, 2017) in order to regulate nest temperatures, but as they reach thermal tolerance limits, they must also undertake more frequent (Clauser and McRae, 2017) or longer incubation recesses (Bueno-Enciso *et al.*, 2017), and may ultimately abandon their nests (Clauser and McRae, 2017; Sharpe *et al.*, 2019).

Here we present the first study of avian reproduction combining both direct observations of incubation behaviour under natural conditions and non-invasive physiological measurements from the same individuals at the same time. We investigated climate effects on the behaviour and physiology of incubating adults in southern pied babblers *Turdoides bicolor* (hereafter ‘pied babblers’), a cooperatively breeding bird. Cooperative species may respond differently to environmental variability compared to pair-breeding or solitary species, because reproductive investment and nest outcomes can be influenced by the presence of helpers (Wiley and Ridley, 2016; van de Ven *et al.*, 2019a), and so we also considered the influence of the number of adults present in each group and checked for interactions between group size and climate (Rubenstein and Lovette, 2007). We hypothesised that high T_air_ would reduce hatching rates via thermoregulatory costs affecting the ability of adult birds to consistently incubate eggs, thus increasing the risk of lethal heat exposure for developing embryos. We addressed this hypothesis by testing predictions related to a) nest outcomes (lower probability of hatching at high T_air_); b) incubation behaviour (reduction in the proportion of time nests are attended at high T_air_); c) the temperatures of unattended nests at high T_air_ (exceeding lethal limits for avian embryos, explaining why hot nests are less likely to hatch); and d) physiological costs of incubation for adults (higher costs of incubation at higher T_air_ evident in patterns of energy expenditure, water balance, and body mass maintenance). We tested part of the latter prediction using a novel, non-invasive doubly-labelled water technique (Anava *et al.*, 2000; Bourne *et al.*, 2019). We further expected that higher rainfall and larger group sizes would be associated with reduced costs and improved nest outcomes in our semi-arid study system.

## Materials and Methods

### Study site and system

Fieldwork took place at the 33km^2^ Kuruman River Reserve (KRR; 26°58’S, 21°49’E) in the southern African Kalahari. Mean summer daily maximum temperatures in the region averaged 34.7 ± 9.7°C and mean annual precipitation averaged 186.2 ± 87.5mm (1995-2015, van de Ven, McKechnie & Cunningham 2019). Rainfall has been declining and high temperature extremes increasing in both frequency and severity over the last 20 years (Kruger and Sekele, 2013; van Wilgen *et al.*, 2016; van de Ven, 2017).

Pied babblers are medium-sized (60–90 g), cooperatively-breeding passerines that live in groups ranging in size from 3–15 adults (Raihani and Ridley, 2007) and are endemic to the region (Ridley, 2016). Resident, territorial groups consist of a single breeding pair (one dominant male and one dominant female) with subordinate helpers of both sexes (Nelson-Flower *et al.*, 2011) and can be reliably located by visits to each territory (Ridley, 2016). Individuals in the study population are habituated to observation by humans at distances of 1–5 m (Ridley and Raihani, 2007), and are individually identifiable by a unique combination of metal and colour leg rings.

Pied babblers build open cup nests, usually in camelthorn *Vachellia erioloba* trees, and usually breed during summer (Ridley, 2016). During each breeding attempt, a clutch of ~3 eggs is laid and incubated for 13–15 days (Ridley and Raihani, 2008). While only the dominant female incubates overnight (Ridley, 2016), during the day all adult group members (individuals > 1 year old), including subordinates, take turns to incubate and the nest is rarely left unattended for more than a few minutes at at time (Ridley and Raihani, 2007; Ridley and van den Heuvel, 2012).

### Data collection

Data were collected during each austral summer breeding season between September 2016 and February 2019 (three breeding seasons in total). We noted daily maximum air temperature (T_max_, in °C) for each observation day and total rainfall in the two months prior to each observation day (mm) using an on-site weather station (Vantage Pro2, Davis Instruments, Hayward, USA), and recorded group size (number of adults) during each breeding attempt. For analyses of nest outcomes, we additionally calculated average T_max_ between initiation of incubation and hatching (Mean T_maxInc_). T_max_ ranged from 20.7°C to 40.8°C (mean = 34.1 ± 4.46), rainfall from 0.2 to 140.2 mm (mean = 15.1 ± 25.1), and group size from 3 to 6 adults (mean = 4 ± 1).

#### Nest outcomes

Monitoring of nest outcomes (*n* = 99 breeding attempts) followed Ridley and van den Heuvel (2012): breeding attempts were defined as discrete clutches laid and incubated; nests were located by observing nest building during weekly monitoring visits; once located, nests were checked approximately every two days to identify incubation start and hatch dates; and nests were categorised as hatched when adult group members were observed carrying food items to the nest, and as failed when nests were left unattended for longer than 90 min on two consecutive monitoring visits or the group was observed building a new nest.

#### Incubation behaviour

Incubation bout and recess data were collected by waiting near the nest at dawn, observing the first bird to replace the dominant female in the morning (05h00–06h48), and remaining with the group all day until 19h00 (*n* = 45 observation days at 35 nests), recording the start and end time of each incubation bout and the duration of any time periods during which the nest was left unattended (recesses). These data were used to calculate the proportion of time per day that clutches were incubated (sum of all incubation bout durations per day / total observation time). Both members of the dominant pair incubated on every observation day, with the help of at least one subordinate group member on most (91%) days.

#### Nest temperatures

We measured operative temperature [T_e_: a measure of thermal load experienced by the bird (Bakken *et al.*, 1985)] using blackbulbs placed in 23 nests within five days of fledge / fail (Griffith *et al.*, 2016), recording constantly for approximately two weeks (13 ± 3 days; range: 10–21 days, *n* = 21,872 T_e_ records in total). Blackbulbs comprised two copper half-spheres (42 mm diameter, which approximates pied babbler thoracic cavity dimensions, and 0.8 mm thick), sealed together using cryanoacrylate adhesive, painted matt black (Carroll *et al.*, 2015; van de Ven *et al.*, 2019b), and containing internally mounted temperature loggers (Thermocron iButton, DS1923, Maxim, Sunnyvale, CA, USA, resolution 0.0625°C) logging at 10-minute intervals (Cunningham *et al.*, 2015; van de Ven *et al.*, 2019b), synchronised with T_air_ records from the onsite weather station. Blackbulbs do not provide a complete representation of thermal conditions experienced by incubating pied babblers because they mimic neither feather arrangement nor colour (Carroll *et al.*, 2017) and do not account for humidity or evaporative heat loss (Bakken *et al.*, 1985). Nonetheless, they provide a relative measure of differences in temperature across nest microsites which cannot be approximated by T_air_ alone (Cunningham *et al.*, 2015).

#### Energy expenditure and water balance

During observation days on which total incubation time for the clutch was recorded, we also obtained detailed physiology [daily energy expenditure (DEE) and water balance] and behaviour (incubation effort) data for a subset of adult birds from the incubating groups (up to four individuals per observation day; mean = 1.6 ± 0.9; *n* = 70 individuals in total). We obtained physiology data from individuals across a range of T_max_ values [*n* = 35 measured on hot days, T_max_ ≥ 35.5°C, identified as a critical temperature threshold in pied babblers (du Plessis *et al.*, 2012; Wiley and Ridley, 2016); *n* = 35 on cool days, T_max_ < 35.5°C] and group sizes (3-6 adults), as well as both sexes (*n* = 38 females, 31 males, 1 unknown sex) and ranks (*n* = 40 dominant birds, 30 subordinate birds). Data on DEE (kJ g^−1^ day^−1^) and water balance were collected using a non-invasive doubly-labelled water (DLW) technique (Anava *et al.*, 2000; Scantlebury *et al.*, 2014), recently validated and described in detail for pied babblers (Bourne *et al.*, 2019).

In brief, selected individuals were dosed with ~50μL of DLW – a non-toxic isotopic solution enriched with oxygen-18 (measured as δ^18^O) and deuterium (measured as δ^2^H) – injected into beetle larvae *Zophobias morio* and fed to the birds between 06h00 and 09h00 on the observation day. Body water samples were then obtained during all daylight hours over a 24 hr observation period by collecting droppings from dosed individuals as they were excreted naturally onto the ground. Water samples were extracted from droppings by cryogenic distillation, using a technique adapted from Priyadarshni *et al.* (2016), and analysed in a PAL autosampler and DLT-100 liquid water isotope analyser (Los Gatos Research, Mountain View, CA, USA) following the procedures described by Smit & McKechnie (2015) and Bourne *et al.* (2019). We calculated CO_2_ production (*r*CO_2_) from the body water pool and the rate of decline of the natural log of the ratio of δ^18^O/δ^2^H (Nagy and Costa, 1980; Speakman, 1997). We used Speakman’s (1997) Equation 17.7 (see eq. 1 below) for calculations of *r*CO_2_ in mol d^−1^ because empirical testing has shown this equation to be the most accurate (Visser *et al.*, 2000) and based on the most realistic assumptions of fractionation during evaporation (Butler *et al.*, 2004; Speakman and Hambly, 2016):

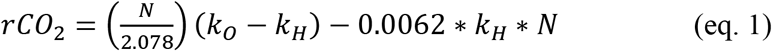

where *N* is moles of body water and values of *k* represent turnover of an isotope identified by the subscript. The divisor of *N* (2.078) accounts for the fact that each molecule of CO_2_ expired removes two molecules of oxygen from the pool and, with the inclusion of the last term (0.0062 • *k*_H_ • *N*), reflects a correction for fractionation. We calculated *k*H in the final term of eq. 1 based on change in ln(*δ*^2^H) between maximally-enriched samples collected at early time points and final samples, where *t* is time (in days) elapsed between early and final samples:

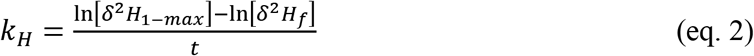

Values of (*k*_*O*_ − *k*_*H*_) can be calculated from the rate of decline of 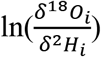, (Nagy and Costa, 1980; Speakman, 1997):

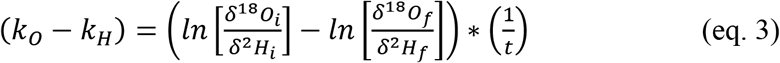

where *δ*^18^O_i_ and *δ*^2^H_i_ are the initial *δ*^18^O and *δ*^2^H values in faeces or blood, and *δ*^18^O_f_ and *δ*^2^H_f_ are the final *δ*^18^O and *δ*^2^H values. *r*CO_2_ was converted from mol d^−1^ to L d^−1^ using the conversion factor 22.4 L of ideal gas per mol at standard temperature and pressure, and L CO_2_ d^−1^ was converted to kJ d^−1^ using the relationship 27.42 kJ/l CO_2_ for an insectivorous bird (Gessaman and Nagy, 1988) and used to estimate DEE (otherwise known as Field Metabolic Rate, in kJ g^−1^ d^−1^).

Water balance was calculated by dividing water influx by water efflux, where values > 1 indicate positive water balance (a hydrated status) and values < 1 indicate negative water balance (a dehydrated status). We used Nagy and Costa’s (1980) Equation 4 (see Equation 4 below) and Equation 6 (see Equation 5 below) to calculate water efflux and water influx (ml H2O kg^−1^d^−1^) respectively:

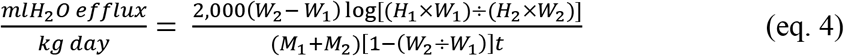

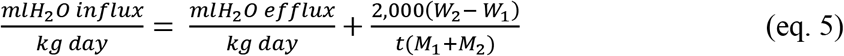

where the subscripts 1 and 2 represent initial and final values respectively, *H* = measured deuterium enrichment levels, *M* = body mass in grams, *W* = the body water pool, and *t* = time in days between initial and final sampling of deuterium enrichment levels. The body water pool was estimated as 69.3% of body mass, as per measured total body water in pied babblers (Bourne *et al.*, 2019).

To identify the proportion of time adult birds dosed with DLW allocated to incubation, we used data collected during ~ 4 x 20-minute continuous time-activity focal behaviour observations (‘focals’, Altmann, 1974) within each of 6 focal sessions per day (mean = 23 focals per bird per day; range: 15–27; *n* = 48 focal days; data were collected from two birds on the same day on 5 occasions, i.e. 10 of the focal days). Focal sessions lasted two hours each, with the first starting at 07h00 and the last at 17h00, and the data were captured on an Android smartphone (Mobicel Trendy), using Prim8 software (McDonald & Johnson, 2014) in which the duration of each observed behaviour is recorded to the nearest second.

#### Body mass

Body mass data were collected from as many adult group members as possible on observation days. These data were obtained by enticing individuals to stand on a top pan balance in exchange for a small food reward (Ridley, 2016), and were collected at dawn on the morning of each observation day (Mass_1_) and again at dawn the following morning (Mass_2_). Body mass change (ΔMb) was calculated in grams as Mass_2_ - Mass_1_ [*n* = 129; pied babblers are size monomorphic (Ridley, 2016) and individuals in the study had similar starting weights (coefficient of variation = 0.07), so using a relative measure (Mass_2_ - Mass_1_ / Mass_1_) did not change interpretation of the models].

### Statistical analyses

Statistical analyses were conducted in the R statistical environment, v 3.6.0 (2017), using the package *lme4* (Bates *et al.*, 2015). Model checking and model selection followed Harrison et al (2018): all continuous explanatory variables were scaled by centering and standardising by the mean; all explanatory variables were tested for correlation with one another and correlated variables (VIF > 2) were not included in the same additive models; Akaike’s information criterion corrected for small sample size (AIC_c_) with maximum likelihood estimation was used to determine which models best explained patterns of variation in the data; model estimates with confidence intervals that did not intersect zero were considered to explain significant patterns within our data; and model fits were evaluated using Normal Q-Q plots, histograms of residuals, and dispersion parameters as appropriate. Rainfall in the two months prior to breeding attempts and breeding season were correlated (*F2,67* = 10.994, *p* < 0.001); we chose the categorical variable ‘breeding season’ for all analyses due to high rainfall occurring in only one breeding season. Quadratic terms for continuous predictors were included when there was no significant linear effect and visualisation of the data suggested a non-linear relationship. Where several models were within 2 AICc of the top model, top model sets were averaged using the package *MuMin* (Barton, 2015). Sensitivity power analysis (Greenland *et al.*, 2016; Champely *et al.*, 2018) suggested sufficient sample size to detect all main effects, but limited power to detect interactions given our data (Table S1). Unless otherwise indicated, summary statistics are presented as means ± one standard deviation.

To determine which variables predicted a) nest outcomes and b) the overall proportion of time that clutches were incubated per day, we used generalised linear mixed-effects models (GLMM) with binomial error structure and logit link function including season, temperature [for a) Mean T_maxInc_; for b) T_max_ on observation day], group size, group size^2, and the interactions between season and group size, and T_max_ and group size, as fixed factors, nest identity as a random factor, and, for b) only, an observation-level random factor to address overdispersion (Harrison, 2014).

To determine which variables predicted DEE (*n* = 68) and water balance (*n* = 69), we used maximum likelihood linear mixed-effects models (LMMs) including season, T_max_, group size, sex, rank, and the interactions between season and group size, and T_max_ and group size, as fixed factors, and bird identity as a random factor. For individuals for which we collected both behaviour and physiology data on the same day, we further considered the influence of proportion of time spent incubating on DEE (*n* = 38) and water balance (*n* = 39), fitting separate linear regressions for hot (≥ 35.5°C) and cool (< 35.5°C) days.

To determine which variables predicted ΔMb, we used the package *segmented* (Muggeo, 2008) to identify the temperature threshold above which ability to maintain body mass between days was compromised, followed by separate LMMs for the data above and below the identified threshold including season, T_max_, group size, sex, rank, and the interactions between season and group size, and T_max_ and group size, as fixed factors, and nest identity as a random factor.

## Results

### Nest outcomes

Of 99 nests monitored over three breeding seasons, 61 hatched and 38 failed. Mean T_max_Inc was the most parsimonious predictor of variation in hatching success in pied babblers (model weight = 0.833). Hatching probability decreased as Mean T_max_Inc increased (Est = −0.303 ± 0.079, 95% CI: −0.467, −0.157, *z* = −3.853; Fig. 1; see Supporting Information Table S2 for full model ouputs) and nests were half as likely to hatch at Mean T_max_Inc > 35.3°C than at cooler incubation temperatures (Fig. 1).

**Figure 1:**
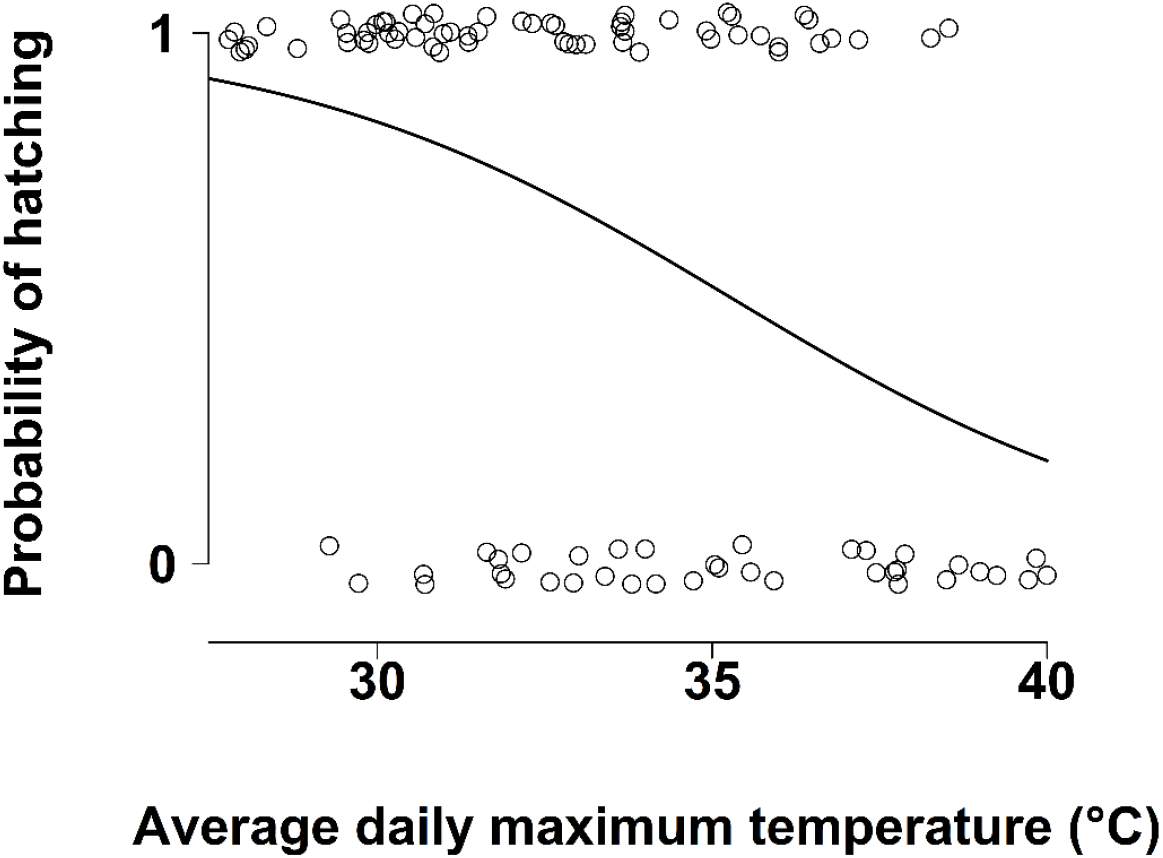
Nest outcomes as a function of mean daily maximum temperatures during incubation. Data from 99 nests by 23 southern pied babbler Turdoides bicolor groups over 3 breeding seasons. Data points are integers (0,1) jittered for improved visibility.

### Nest attendance

The proportion of time between dawn and 19h00 that clutches were incubated ranged from 57.3 to 100% (mean = 95.1 ± 8.2%). Only three nests were incubated for < 80% of daylight hours, all of which were observed on days with T_max_ > 37°C and all of which failed. Temperature (T_max_) was the most parsimonious predictor of variation in incubation time (model weight = 0.498): incubation declined as temperatures increased (Est = −1.588 ± 0.466, 95% CI: −2.528, −0.648, *z* = −3.311; Fig. 2; see Supporting Information Table S3 for full model ouputs).

**Figure 2:**
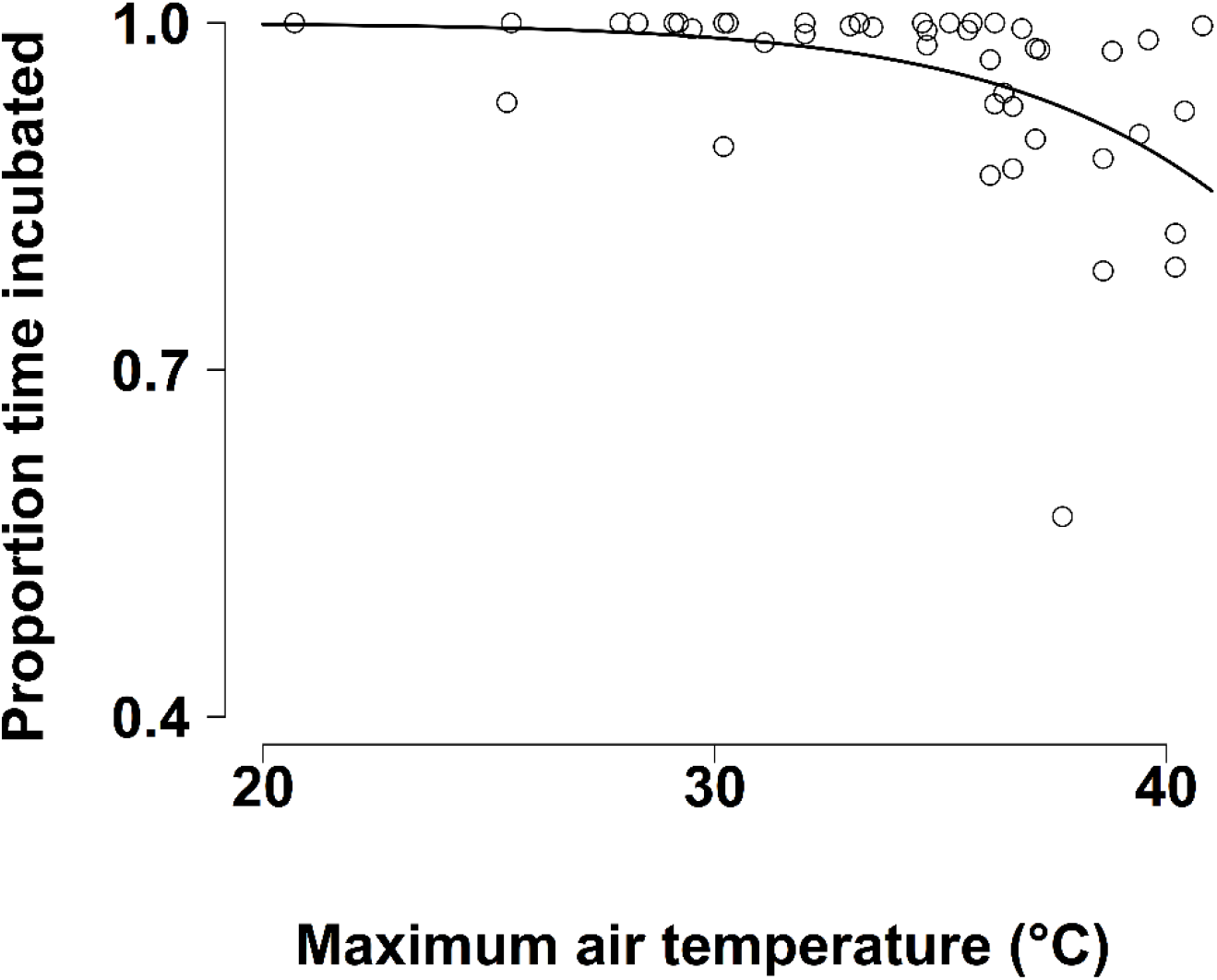
Proportion of time that the clutch was incubated as a function of maximum air temperature on the observation day. Data from 46 observation days at 35 southern pied babbler Turdoides bicolor nests over 3 breeding seasons.

### Nest temperatures

Diurnal nest T_e_ always exceeded T_air_ (06h00–19h00; mean difference = 7.9 ± 11.2°C, range: 0.01–31.8°C; Fig. 3a; Table S7). At the coolest T_air_ recorded during the day (~8°C, *n* = 2 days), nest T_e_ averaged 10.1 ± 0.7 °C (range: 8.8–11.6°C; *n* = 5 nests), and at the warmest T_air_ recorded during the day (~41°C, *n* = 1 day), nest T_e_ averaged 44.4 ± 2.8°C (range: 40.9–49.1°C; *n* = 1 nest). Nest T_e_ increased significantly with T_air_ (linear regression; Est = 1.207 ± 0.005, 95% CI: 1.196, 1.217, *t* = 229.2; Fig. 3b). The highest nest T_e_ recorded was 65°C and operative temperatures > 60°C were recorded at two nests for T_air_ between ~30°C and ~37°C. We recorded 2,379 instances of T_e_ in unattended nests > 41°C (10.8% of all T_e_ records, 22 of 23 nests; mean = 108.1 ± 84.6 instances per nest, range: 30-295), identified as a potentially lethal temperature for avian embryos (Webb, 1987; DuRant *et al.*, 2013), and 487 instances of T_e_ in unattended nests >50°C (2.2% of all T_e_ records, 17 of 23 nests; mean = 28.6 ± 41.4 instances per nest, range: 1-163), lethal for many arid zone species (Grant, 1982; Reyna and Burggren, 2012; Griffith *et al.*, 2016).

**Figure 3:**
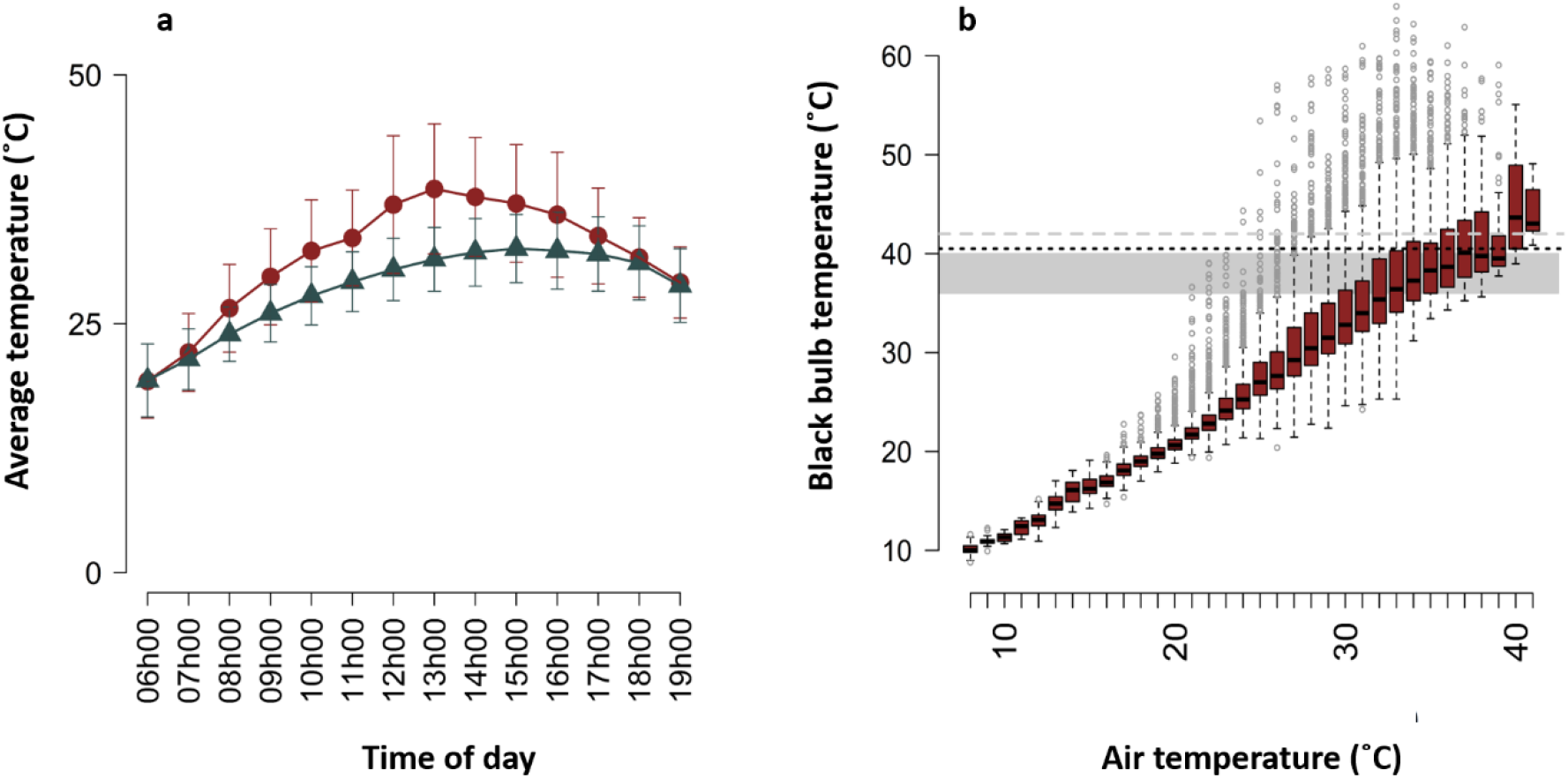
(a) Comparison of average temperatures recorded per hour between 06h00 and 19h00 (mean ± sd) by an onsite weather station (blue triangles) and blackbulbs placed in vacated southern pied babbler Turdoides bicolor nests (red circles); (b) blackbulb temperature as a function of air temperature. Boxplots show the median and interquartile range (IQR) of operative temperature for each air temperature value rounded to the nearest digit. Whiskers indicate the lowest and highest value datapoints within 1.5*IQR. Points plotted beyond the whiskers represent a relatively small number of extreme values in this large dataset, n = 21,872 temperature records. The optimal temperature range for avian embryo development (36–40°C, shaded area), the lowest potential lethal temperature for avian embryos given prolonged exposure (40.5°C, black dotted line), and the average upper critical limit for thermoneutrality in passerines (41°C, grey dashed line) are indicated on (b).

### Energy expenditure and water balance

We quantified DEE (*n* = 68; mean = 1.613 ± 0.463 kJ^−1^g^−1^d, range: 0.639-2.855 kJ^−1^g^−1^d) and water balance (*n* = 69; mean = 1.034 ± 0.116, range: 0.869-1.691; where 1 = neutral water balance) in birds from incubating groups. Temperature (T_max_) was the most parsimonious predictor of variation in DEE (model weight = 0.553): DEE declined as temperature increased (Est = −0.223 ± 0.046, 95% CI: −0.315, −0.131, *z* = 4.762; Fig. 4; see Supporting Information Table S8 for full model ouput). Variation in water balance was not predicted by any of the variables included in our models (Table S9). Our within-individual physiology and behaviour data showed no evidence that DEE was predicted by the proportion of time spent incubating on either hot or cool days (*n* = 38; Fig.5a, Table 1). However, these data showed that pied babblers’ ability to maintain neutral or positive water balance declined with an increasing proportion of time spent incubating on hot days, but not on cool days (*n* = 39; Fig.5b, Table 1).

**Figure 2:**
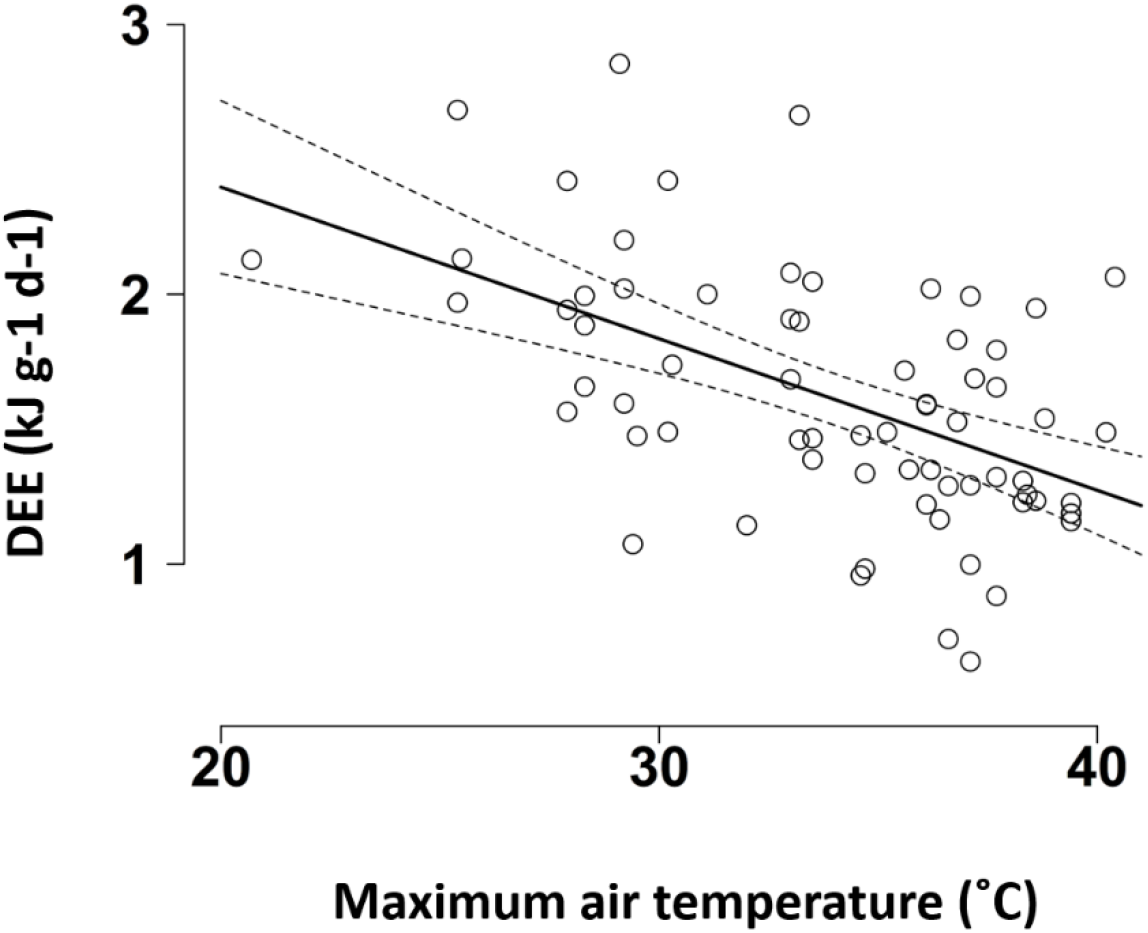
Variation in daily energy expenditure by maximum air temperature (°C) on the measurement day in southern pied babblers Turdoides bicolor

**Figure 3:**
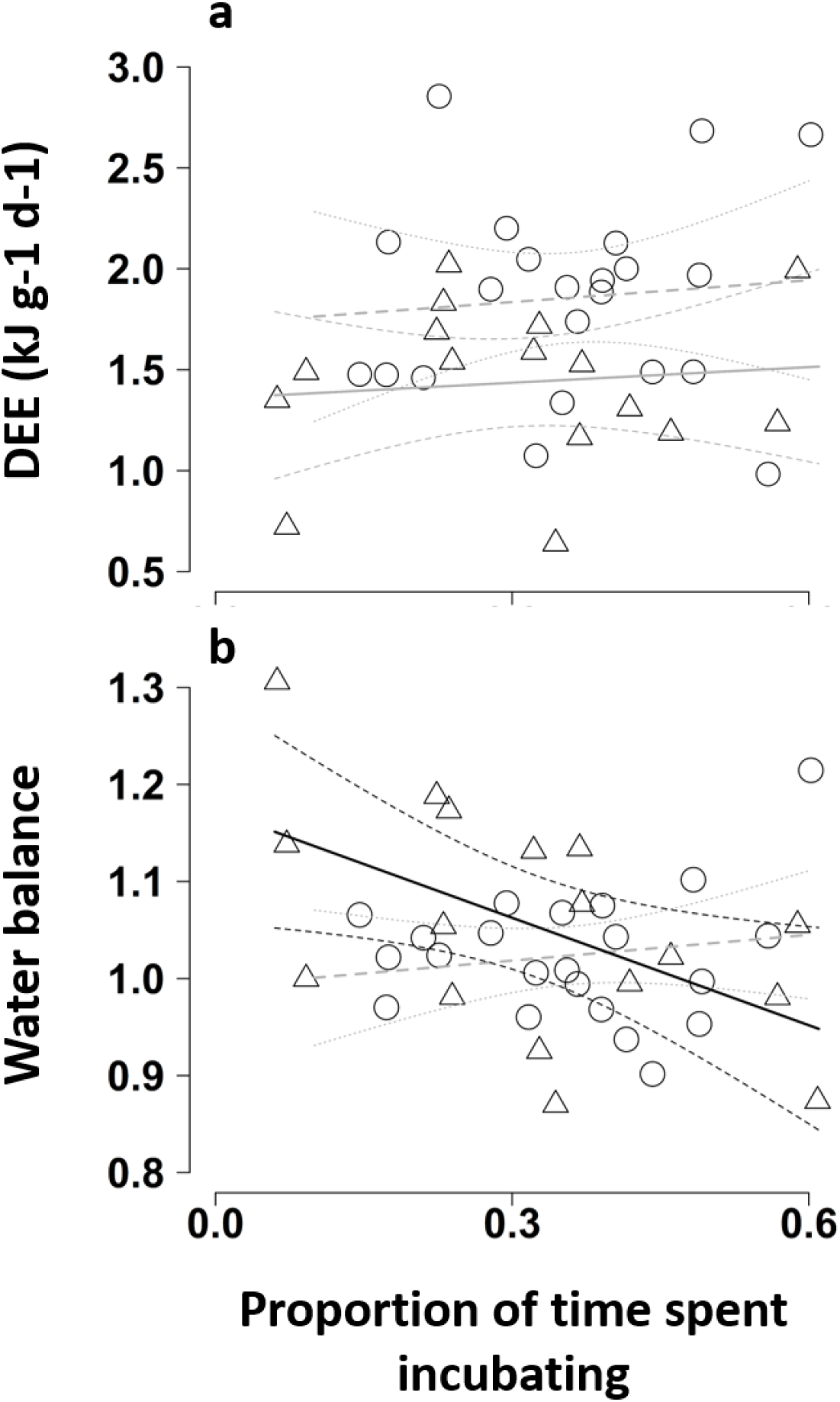
Influence of proportion of time southern pied babblers Turdoides bicolor spent incubating on cool (T_max_ < 35.5°C, open circles, dashed lines, dotted 95% CIs) and hot (T_max_ ≥ 35.5°C, open triangles, solid lines, dashed 95% CIs) days on the (a) daily energy expenditure and (b) water balance of incubating birds. Model fit lines for non-significant relationships are faded to grey.

**Table 1:** Daily energy expenditure and water balance as a function of proportion of time spent incubating, analysed separately for cool (T_max_ < 35.5°C) and hot (T_max_ ≥ 35.5°C) days. Significant relationships are shown in bold.

| Response | *n* | Temperature | Estimate | Std error | 95% CI | *t* value | *p* value |
| --- | --- | --- | --- | --- | --- | --- | --- |
| Daily energy expenditure | 22 | Cool | 0.360 | 0.871 | -1.456/2.177 | 0.414 | 0.684 |
|  | 16 | Hot | 0.258 | 0.662 | -1.162/1.678 | 0.390 | 0.703 |
| Water balance | 22 | Cool | 0.089 | 0.117 | -0.155/0.332 | 0.758 | 0.457 |
|  | 17 | Hot | **-0.369** | **0.149** | **-0.687/-0.052** | **-2.480** | **0.026** |

### Body mass

Mass change over 24hrs averaged 0.29 ± 2.26 g (range: −4.3-6.3 g, or −9.1-8.5% of body mass; *n* = 120 individuals). We detected a threshold T_max_ at 36.2°C (95% CI: 34.1, 38.2°C). At T_max_ < 36.2°C (*n* = 72), ΔMb was not predicted by any of the independent variables included (Table S10). At T_max_ ≥ 36.2°C (*n* = 48), T_max_ was the only predictor that significantly influenced ΔM_b_ (model weight = 0.633), with mass loss becoming more likely as temperatures increased (Est = −0.926 ± 0.318, 95% CI: −1.609, −0.303, *t* = −2.908; Fig. 6; see Supporting Information Table S11 for full model outputs).

**Figure 6:**
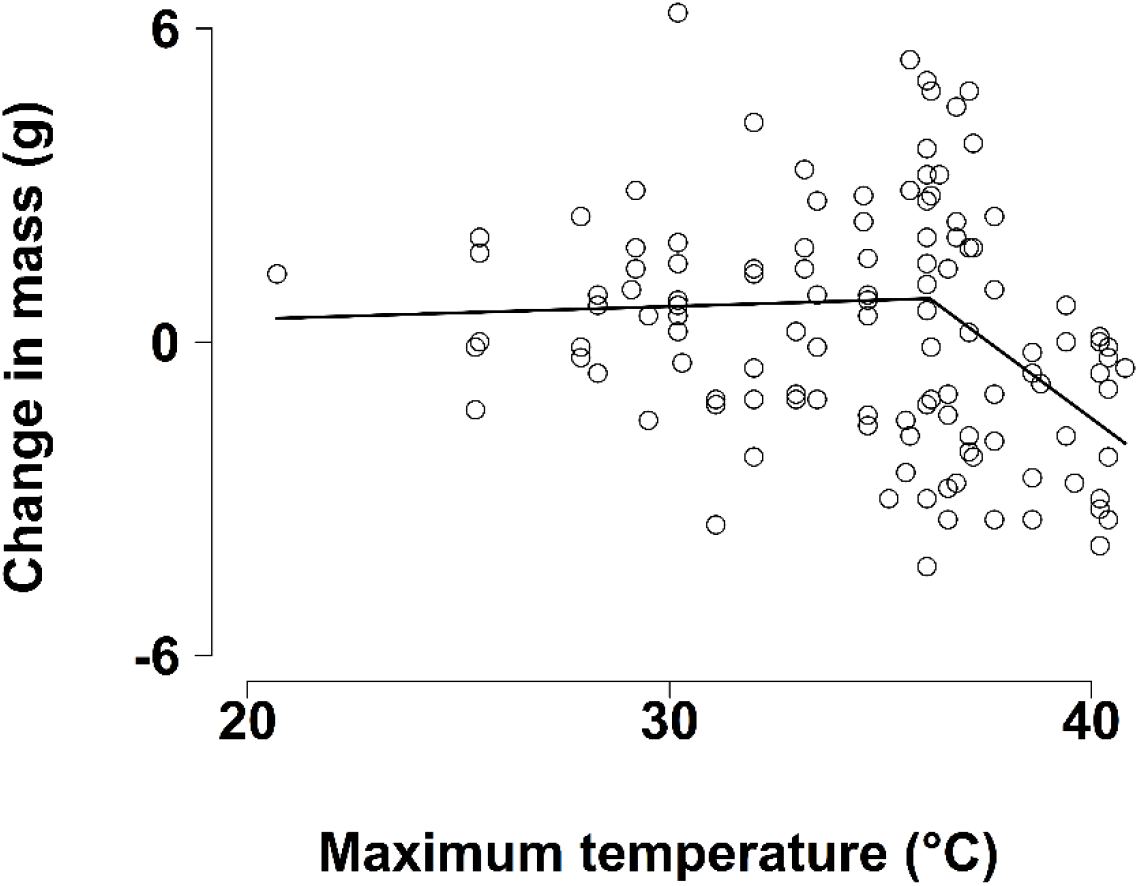
Change in southern pied babbler Turdoides bicolor body mass (g) from one morning to the next as a function of maximum air temperature (°C) on the observation day. Line represents the segmented linear regressions for the relationship between mass change and temperature above and below 36.2°C, i.e. no relationship below the threshold temperature and a significant negative relationship above the temperature threshold.

## Discussion

Pied babblers exhibited poor hatching success at high temperatures. Employing a novel combination of non-invasive DLW and field-based behaviour observations, we demonstrated that a) pied babblers generally incubated their nests almost constantly (95% of daylight hours), but the proportion of time that nests were attended declined with increasing T_air_ (Bueno-Enciso *et al.*, 2017; Clauser and McRae, 2017); 2) T_e_ in unattended nests was substantially higher than T_air_ and frequently exceeded widely reported lethal limits for avian embryos (Webb, 1987; Conway and Martin, 2000; Birkhead *et al.*, 2008; DuRant *et al.*, 2013; Wada *et al.*, 2015) and the inflection T_air_ values above which passerine birds rapidly increase rates of evaporative water loss via panting (McKechnie *et al.*, 2017; Smith *et al.*, 2017); c) pied babblers incurred water costs, but not energy costs, of incubation at high temperatures (Smit and McKechnie, 2015; Cooper *et al.*, 2019); and d) pied babblers from incubating groups lost mass during very hot weather (du Plessis *et al.*, 2012; Sharpe *et al.*, 2019; van de Ven *et al.*, 2019b). Multiple lines of evidence suggest that, during very hot periods, incubating pied babblers leave nests unattended to avoid dehydration (Downs and Ward, 1997; Clauser and McRae, 2017), rather than to take advantage of ambient incubation (Londoño *et al.*, 2008; Boulton *et al.*, 2010; Bambini *et al.*, 2019). With T_e_ in unattended nests regularly exceeding lethal limits for avian embryos, reduced nest attendance at high T_air_ may contribute to reduced hatching success during hot incubation periods.

Our finding that incubating pied babblers failed to maintain water balance when incubating for long periods of time on hot days, but not on cool days, is novel and strongly suggests that birds incubating at high temperatures leave the nest as a result of the water costs incurred. This contrasts with commonly held views that birds should benefit from higher tempertures during incubation because of the opportunity to leave the nest without compromising embryo development as a result of eggs cooling (Conway and Martin, 2000; Bambini *et al.*, 2019). Incubating birds cannot fully engage in normal behavioural thermoregulation, such as retreating to the shade or adjusting foraging and drinking behaviours (Smit *et al.*, 2016; Abdu *et al.*, 2018; Cooper *et al.*, 2019), and rely on evaporative cooling to maintain body temperature below lethal levels (Grant, 1982; Brown and Downs, 2003; O’Connor *et al.*, 2018), presumably at high water cost to themselves given high nest T_e_. Lethal dehydration has resulted in mass mortality of birds (McKechnie and Wolf, 2010; Gardner *et al.*, 2019) and mammals (Welbergen *et al.*, 2008; Ratnayake *et al.*, 2019) and the water turnover rates of birds in arid environments tend to be frugal (Williams and Tieleman, 2005; Cooper *et al.*, 2019). That temperature did not affect water balance except in interaction with the proportion of time spent incubating, provides an indication of just how important it is for birds to maintain water balance over short timescales in hot and dry environments.

Pied babblers build open cup nests in sparse vegetation (Ridley, 2016), and the T_e_ we recorded in unattended pied babbler nests regularly exceeded a) temperatures at which evaporative water loss increases rapidly in passerine birds (41°C, McKechnie et al., 2017; Smith et al., 2017), b) optimal temperatures for embryo development in passerines (36–40°C, DuRant et al., 2013), and c) lethal temperature limits for developing avian embryos (40.5°C-51°C, DuRant et al., 2013; Grant, 1982; Griffith et al., 2016; Stoleson & Beissinger, 1999; Webb, 1987). Such high nest temperatures have been recorded in several bird species nesting in exposed sites (Brown and Downs, 2003; Tieleman *et al.*, 2008; AlRashidi *et al.*, 2011; Clauser and McRae, 2017). While some arid zone species exhibit high heat tolerance in developing embryos (Grant, 1982; Reyna and Burggren, 2012; Griffith *et al.*, 2016), leaving nests unattended for long periods of time during the heat of the day exposes developing avian embryos to high temperatures (Mayer *et al.*, 2009; Carroll *et al.*, 2015; DuRant *et al.*, 2019), potentially exceeding lethal limits (Webb, 1987) and risking embryo death (Birkhead *et al.*, 2008; Wada *et al.*, 2015; Clauser and McRae, 2017) or other problems, such as increased hatching asynchrony (Griffith *et al.*, 2016). It is therefore likely that near-constant incubation and / or shading is both highly desirable (Grant, 1982), in order to limit exposure of embryos to excessive heat, and also difficult to sustain at high temperatures, because birds prevent body temperature exceeding lethal limits by evaporative cooling (Albright *et al.*, 2017; O’Connor *et al.*, 2017; McKechnie and Wolf, 2019). The reduced nest attendance we observed at high temperatures likely indicates that parental investment in incubation is constrained by the water costs of heat exposure (Amat and Masero, 2004; Coe *et al.*, 2015), and may suggest progress towards eventual nest abandonment (Stoleson and Beissinger, 1999; Sharpe *et al.*, 2019).

## Conclusions

Given that a) pied babblers incubate their eggs almost constantly during the day, b) instances where lower than normal incubation constancy was observed all occurred on hot days, and c) unusually low incubation constancy was followed by nest abandonment or failure, we suggest that reduced incubation constancy at high temperatures contributes to hatching failure by increasing the risk of exposure of embryos to lethal temperatures. We cannot directly test for causal relationships between effects of temperature on the behaviour and physiology of incubating pied babblers and hatching success, which would require an experimental approach or at least observations over multiple days within the same breeding attempts. However, we present multiple lines of evidence suggesting that pied babblers should stay on their nests and avoid ambient incubation at high temperatures, to prevent embryos from overheating, but may be constrained from doing so by the thermoregulatory costs of incubation at high temperatures.

Rather than strategically leaving the nest to take advantage of opportunities for ambient incubation, pied babblers appear to be leaving their nests on hot afternoons due to the water costs incurred as a result of incubating at high temperatures. We argue that considering both behaviour and physiology simultaneously in the same individuals, at the same time, under natural conditions, provides invaluable insights into the thermal constraints under which incubating birds operate. As we found no effect of group size on the responses we measured, either alone or in interaction with environmental factors, we further suggest that cooperative breeding may not confer an advantage over non-cooperative breeding strategies in buffering against hot weather during the incubation phase.

Although parental care strategies are flexible in response to both climate and social conditions (Clutton-Brock *et al.*, 2004; Langmore *et al.*, 2016), these strategies have limits (Clauser and McRae, 2017; Sharpe *et al.*, 2019). Given that both mean temperatures and hot extremes are increasing in frequency under global climate change (IPCC, 2013), the incubation period could become a major bottleneck for reproduction across species with different reproductive strategies. Birds will likely incur ever greater thermoregulatory costs of incubation as temperatures rise, leading to reduced nest attendance, potential overheating of eggs, and ultimately, compromised population replacement and persistence.

## Acknowledgements

We thank the management teams at the Kuruman River Reserve (KRR) and surrounding farms, Van Zylsrus, South Africa, for making the work possible, and Sello Matjee, Paige Ezzey, and Lesedi Moagi for fieldwork assistance. The KRR was financed by the Universities of Cambridge and Zurich, the MAVA Foundation, and the European Research Council (Grant No. 294494 to Tim Clutton-Brock) and received logistical support from the Mammal Research Institute of the University of Pretoria. This work was supported by the Australian Research Council (FT110100188 to ARR); the BBSRC David Phillips Fellowship (BB/J014109/1 to CNS); the British Ornithologists’ Union; the DST-NRF Centre of Excellence at the FitzPatrick Institute for African Ornithology; the Oppenheimer Memorial Trust (20747/01 to ARB); the University of Cape Town; and the National Research Foundation of South Africa (Grant No. 110506 to AEM, Grant No. 99050 and 118627 to SJC). The opinions, findings and conclusions are those of the authors alone, and the National Research Foundation accepts no liability whatsoever in this regard.

## Literature cited

Abdu S, McKechnie AE, Lee ATK, Cunningham SJ (2018) Can providing shade at water points help Kalahari birds beat the heat? J Arid Environ 152: 21–27.

Albright TP, Mutiibwa D, Gerson AR, Smith EK, Talbot WA, O’Neill JJ, McKechnie AE, Wolf BO (2017) Mapping evaporative water loss in desert passerines reveals an expanding threat of lethal dehydration. Proc Natl Acad Sci 114: 2283–2288.

AlRashidi M, Kosztolányi A, Shobrak M, Küpper C, Székely T (2011) Parental cooperation in an extreme hot environment: natural behaviour and experimental evidence. Anim Behav 82: 235–243.

Altmann J (1974) Observational study of behaviour: sampling methods. Behaviour 49: 227–266.

Amat JA, Masero JA (2004) How Kentish plovers, Charadrius alexandrinus, cope with heat stress during incubation. Behav Ecol Sociobiol 56: 26–33.

Anava A, Kam M, Shkolnik A, Degen AA (2000) Seasonal field metabolic rate and dietary intake in Arabian Babblers (*Turdoides squamiceps*) inhabiting extreme deserts. Funct Ecol 14: 607–613.

Ardia DR, Pérez JH, Clotfelter ED (2010) Experimental cooling during incubation leads to reduced innate immunity and body condition in nestling tree swallows. Proc R Soc B Biol Sci 277: 1881–1888.

Bakken GS, Santee WR, Erskine DJ (1985) Operative and standard operative temperature: tools for thermal energetics studies. Am Zool 25: 933–943.

Bambini G, Schlicht E, Kempenaers B (2019) Patterns of female nest attendance and male feeding throughout the incubation period in blue tits *Cyanistes caeruleus*. Ibis (Lond 1859) 161: 50–65.

Barton K (2015) MuMIn: Multi-model inference. https://cran.r-project.org/package=MuMin

Bates D, Maechler M, Bolker B, Walker S (2015) Fitting linear mixed effects models using lme4. J Stat Softw 67: 1–48.

Birkhead TR, Hall J, Schut E, Hemmings N (2008) Unhatched eggs: methods for discriminating between infertility and early embryo mortality. Ibis (Lond 1859) 150: 508–517.

Boulton RL, Richard Y, Armstrong DP (2010) The effect of male incubation feeding, food and temperature on the incubation behaviour of New Zealand robins. Ethology 116: 490–497.

Bourne AR, McKechnie AE, Cunningham SJ, Ridley AR, Woodborne SM, Karasov WH (2019) Non invasive measurement of metabolic rates in wild, free living birds using doubly labelled water. Funct Ecol 33: 162–174.

Brown M, Downs CT (2003) The role of shading behaviour in the thermoregulation of breeding crowned plovers (*Vanellus coronatus*). J Therm Biol 28: 51–58.

Bueno-Enciso J, Barrientos R, Ferrer ES, Sanz JJ (2017) Do extended incubation recesses carry fitness costs in two cavity-nesting birds? J F Ornithol 88: 146–155.

Butler PJ, Green JA, Boyd IL, Speakman JR (2004) Measuring meatabolic rate in the field: the pros and cons of the doubly labeled water and heart rate methods. Funct Ecol 18: 168–183.

Cahill AE, Aiello-Lammens ME, Caitlin Fisher-Reid M, Hua X, Karanewsky CJ, Ryu HY, Sbeglia GC, Spagnolo F, Waldron JB, Warsi O, et al. (2013) How does climate change cause extinction? Proc R Soc B Biol Sci 280: 20121890.

Carroll JM, Davis CA, Elmore RD, Fuhlendorf SD (2015) A ground-nesting galliform’s response to thermal heterogeneity: implications for ground-dwelling Birds. PLoS One 10: 1–21.

Carroll JM, Davis CA, Elmore RD, Fuhlendorf SD (2017) Using a historic drought and high-heat event to validate thermal exposure predictions for ground-dwelling birds. Ecol Evol 7: 6413–6422.

Champely S, Ekstrom C, Dalgaard P, Gill J, Weibelzahl S, Anandkumar A, Ford C, Volcic R, De Rosario H (2018) pwr: Basic functions for power analysis. https://github.com/heliosdrm/pwr

Clauser AJ, McRae SB (2017) Plasticity in incubation behaviour and shading by king rails (Rallus elegans) in response to temperature. J Avian Biol 48: 479–488.

Clutton-Brock TH, Russell AF, Sharpe LL (2004) Behavioural tactics of breeders in cooperative meerkats. Anim Behav 68: 1029–1040.

Coe BH, Beck ML, Chin SY, Jachowski CMB, Hopkins WA (2015) Local variation in weather conditions influences incubation behavior and temperature in a passerine bird. J Avian Biol 46: 385–394.

Conradie S, Woodborne SM, Cunningham SJ, McKechnie AE (2019) Chronic, sublethal effects of high temperatures will cause severe declines in southern African arid-zone birds during the 21st century. Proc Natl Acad Sci 116: 14065–14070.

Conrey RY, Skagen SK, Yackel Adams AA, Panjabi AO (2016) Extremes of heat, drought and precipitation depress reproductive performance in shortgrass prairie passerines. Ibis (Lond 1859) 158: 614–629.

Conway CJ, Martin Te (2000) Effects of ambient temperature on avian incubation behavior. Behav Ecol 11: 178–188.

Cooper CE, Withers PC, Hurley LL, Griffith SC (2019) The field metabolic rate, water turnover, and feeding and drinking behavior of a small avian desert granivore during a summer heatwave. Front Physiol 10: 1405.

Cunningham SJ, Martin RO, Hockey PAR (2015) Can behaviour buffer the impacts of climate change on an arid-zone bird? Ostrich 86: 119–126.

Cunningham SJ, Martin RO, Hojem CL, Hockey PAR (2013) Temperatures in excess of critical thresholds threaten nestling growth and survival in a rapidly-warming arid savanna: a study of common fiscals. PLoS One 8: e74613.

Downs CT, Ward D (1997) Does shading behavior of incubating shorebirds in hot environments cool the eggs or the adults? Auk 114: 717–724.

du Plessis KL, Martin RO, Hockey PAR, Cunningham SJ, Ridley AR (2012) The costs of keeping cool in a warming world: Implications of high temperatures for foraging, thermoregulation and body condition of an arid-zone bird. Glob Chang Biol 18: 3063–3070.

DuRant SE, Hopkins WA, Hepp GR, Walters JR (2013) Ecological, evolutionary, and conservation implications of incubation temperature-dependent phenotypes in birds. Biol Rev 88: 499–509.

DuRant SE, Willson JD, Carroll RB (2019) Parental effects and climate change: will avian incubation behavior shield embryos from increasing environmental temperatures? Integr Comp Biol 59: 1068–1080.

Gardner JL, Amano T, Peters A, Sutherland WJ, Mackey B, Joseph L, Stein J, Ikin K, Little R, Smith J, et al. (2019) Australian songbird body size tracks climate variation: 82 species over 50 years. Proc R Soc B Biol Sci 286: 20192258.

Gessaman JA, Nagy KA (1988) Energy metabolism: errors in gas exchange conversion factors. Physiol Zool 61: 507–513.

Grant GS (1982) Avian incubation: egg temperature, nest humidity, and behavioral thermoregulation in a hot environment. Ornithol Monogr 30: 1–75.

Greenland S, Senn SJ, Rothman KJ, Carlin JB, Poole C, Goodman SN, Altman DG (2016) Statistical tests, P values, confidence intervals, and power: a guide to misinterpretations. Eur J Epidemiol 31: 337–350.

Griffith SC, Mainwaring MC, Sorato E, Beckmann C (2016) High atmospheric temperatures and ‘ambient incubation’ drive embryonic development and lead to earlier hatching in a passerine bird. R Soc Open Sci 3: 150371.

Harrison XA (2014) Using observation-level randomeffects to model overdispersion in count data in ecology and evolution. PeerJ 2014. doi:10.7717/peerj.616

Harrison XA, Donaldson L, Correa-cano ME, Evans J, Fisher DN, Goodwin CED, Robinson BS, Hodgson DJ, Inger R (2018) A brief introduction to mixed effects modelling and multi-model inference in ecology. Peer J 6: 1–32.

Iknayan KJ, Beissinger SR (2018) Collapse of a desert bird community over the past century driven by climate change. Proc Natl Acad Sci 115: 8597–8602.

IPCC (2013) Climate Change 2013: The Intergovernmental Panel on Climate Change Fifth Assessment Report. Cambridge University Press, Cambridge.

Kruger AC, Sekele SS (2013) Trends in extreme temperature indices in South Africa: 1962-2009. Int J Climatol 33: 661–676.

Langmore NE, Bailey LD, Heinsohn RG, Russell AF, Kilner RM (2016) Egg size investment in superb fairy-wrens: helper effects are modulated by climate. Proc R Soc B 283: 10–12.

Londoño GA, Levey DJ, Robinson SK (2008) Effects of temperature and food on incubation behaviour of the northern mockingbird, Mimus polyglottos. Anim Behav 76: 669–677.

Mayer PM, Smith LM, Ford RG, Watterson DC, Mccutchen MD, Ryan MR (2009) Nest construction by a ground-nesting bird represents a potential trade-off between egg crypticity and thermoregulation. Oecologia 159: 893–901.

McDonald M, Johnson S (2014) “There’s an app for that”: a new program for the collection of behavioural field data. Anim Behav 95: 81–87.

McDonald S, Schwanz LE (2018) Thermal parental effects on offspring behaviour and their fitness consequences. Anim Behav 135: 45–55.

McKechnie AE (2019) Physiological and morphological effects of climate change. In: Dunn PO, Moller AP, eds. Effects of Climate Change on Birds. Oxford University Press, Oxford, pp 120–133.

McKechnie AE, Gerson AR, McWhorter TJ, Smith EK, Talbot WA, Wolf BO (2017) Avian thermoregulation in the heat: evaporative cooling in five Australian passerines reveals within-order biogeographic variation in heat tolerance. J Exp Biol 220: 2436–2444.

McKechnie AE, Wolf BO (2010) Climate change increases the likelihood of catastrophic avian mortality events during extreme heat waves. Biol Lett 6: 253–6.

McKechnie AE, Wolf BO (2019) The physiology of heat tolerance in small endotherms. Physiology 34: 302–313.

Mortensen JL, Reed JM (2018) Parental incubation patterns and the effect of group size in a Neotropical cooperative breeder. Auk Ornithol Adv 135: 669–692.

Muggeo VM. (2008) Segmented: an R package to fit regression models with broken-line relationships. R News 8: 20–25.

Nagy KA, Costa DP (1980) Water flux in animals: analysis of potential errors in the tritiated water method. Am J Physiol 238: 454–65.

Nelson-Flower MJ, Hockey PAR, O’Ryan C, Raihani NJ, Du Plessis MA, Ridley AR (2011) Monogamous dominant pairs monopolize reproduction in the cooperatively breeding pied babbler. Behav Ecol 22: 559–565.

Nord A, Cooper CB (2020) Night conditions affect morning incubation behaviour differently across a latitudinal gradient. Ibis (Lond 1859). doi:10.1111/ibi.12804

Nord A, Sandell MI, Nilsson JÅ (2010) Female zebra finches compromise clutch temperature in energetically demanding incubation conditions. Funct Ecol 24: 1031–1036.

Nwaogu CJ, Dietz MW, Tieleman BI, Cresswell W (2017) Breeding limits foraging time: evidence of interrupted foraging response from body mass variation in a tropical environment. J Avian Biol 48: 563–569.

O’Connor RS, Brigham RM, McKechnie AE (2018) Extreme operative temperatures in exposed microsites used by roosting rufous-cheeked nightjar (*Caprimulgus rufigena*): implications for water balance under current and future climate conditions. Can J Zool 96: 1122–1129.

O’Connor RS, Wolf BO, Brigham RM, McKechnie AE (2017) Avian thermoregulation in the heat: efficient evaporative cooling in two southern African nightjars. J Comp Physiol B Biochem Syst Environ Physiol 187: 477–491.

Priyadarshini KVR, Prins HHT, de Bie S, Heitkönig IMA, Woodborne S, Gort G, Kirkman K, Ludwig F, Dawson TE, de Kroon H (2016) Seasonality of hydraulic redistribution by trees to grasses and changes in their water-source use that change tree-grass interactions. Ecohydrology 9: 218–228.

R Core Team (2017) R: A Language and Environment for Statistical Computing. R Foundation for Statistical Computing. R Foundation for Statistical Computing, Vienna.

Raihani NJ, Ridley AR (2007) Variable fledging age according to group size: trade-offs in a cooperatively breeding bird. Biol Lett 3: 624–627.

Ratnayake HU, Kearney MR, Govekar P, Karoly D, Welbergen JA (2019) Forecasting wildlife die-offs from extreme heat events. Anim Conserv 22: 386–395.

Reyna KS, Burggren WW (2012) Upper lethal temperatures of northern bobwhite embryos and the thermal properties of their eggs. Poult Sci 91: 41–46.

Ridley AR (2016) Southern pied babblers: The dynamics of conflict and cooperation in a group-living society. In: Dickinson JL, Koenig W, eds. Cooperative Breeding in Vertebrates: Studies of Ecology, Evolution, and Behavior. Cambridge University Press, Cambridge, pp 115–132.

Ridley AR, Raihani NJ (2007) Variable postfledging care in a cooperative bird: causes and consequences. Behav Ecol 18: 994–1000.

Ridley AR, Raihani NJ (2008) Task partitioning increases reproductive output in a cooperative bird. Behav Ecol 19: 1136–1142.

Ridley AR, van den Heuvel I (2012) Is there a difference in reproductive performance between cooperative and non-cooperative species? A southern African comparison. Behaviour 8: 821–848.

Ripple WJ, Wolf C, Newsome TM, Barnard P, Moomaw WR (2019) World scientists’ warning of a climate emergency: a second notice. Bioscience 67: 1026–1028.

Rosenberg K V., Dokter AM, Blancher PJ, Sauer JR, Smith AC, Smith PA, Stanton JC, Panjabi A, Helft L, Parr M, et al. (2019) Decline of the North American avifauna. Science (80-) 336: 120–124.

Rubenstein DR, Lovette IJ (2007) Temporal environmental variability drives the evolution of cooperative breeding in birds. Curr Biol 17: 1414–1419.

Saino N, Ambrosini R, Rubolini D, Von Hardenberg J, Provenzale A, Hüppop K, Hüppop O, Lehikoinen A, Lehikoinen E, Rainio K, et al. (2011) Climate warming, ecological mismatch at arrival and population decline in migratory birds. Proc R Soc B Biol Sci 278: 835–842.

Scantlebury DM, Mills MGL, Wilson RP, Wilson JW, Mills MEJ, Durant SM, Bennett NC, Bradford P, Marks NJ, Speakman JR (2014) Flexible energetics of cheetah hunting strategies provide resistance against kleptoparasitism. Science (80-) 346: 79–81.

Sharpe L, Cale B, Gardner JL (2019) Weighing the cost: the impact of serial heatwaves on body mass in a small Australian passerine. J Avian Biol 50: jav.02355.

Smit B, McKechnie AE (2015) Water and energy fluxes during summer in an arid-zone passerine bird. Ibis (Lond 1859) 157: 774–786.

Smit B, Zietsman G, Martin RO, Cunningham SJ, Mckechnie AE, Hockey PAR (2016) Behavioural responses to heat in desert birds: implications for predicting vulnerability to climate warming. Clim Chang Responses 3: 1–14.

Smith EK, O’Neill JJ, Gerson AR, McKechnie AE, Wolf BO (2017) Avian thermoregulation in the heat: resting metabolism, evaporative cooling and heat tolerance in Sonoran Desert songbirds. J Exp Biol 220: 3290–3300.

Speakman JR (1997) Doubly Labelled Water: Theory and Practice. Springer Science & Business Media., New York.

Speakman JR, Hambly C (2016) Using doubly-labelled water to measure free-living energy expenditure: some old things to remember and some new things to consider. Comp Biochem Physiol Part A 202: 3–9.

Stevenson IR, Bryant DM (2000) Avian phenology: climate change and constraints on breeding. Nature 406: 366–367.

Stillman JH (2019) Heat waves, the new normal: summertime temperature extremes will impact animals, ecosystems, and human communities. Physiology 34: 86–100.

Stoleson SH, Beissinger SR (1999) Egg viability as a constraint on hatching synchrony at high ambient temperatures. J Anim Ecol 68: 951–962.

Tieleman BI, van Noordwijk JF, Williams JB (2008) Nest site selection in a hot desert: trade-off between microclimate and predation risk. Condor 110: 116–124.

van de Ven TMFN (2017) Implications of Climate Change on the Reproductive Success of the Southern Yellow-Billed Hornbill Tockus Leucomelas. PhD thesis, University of Cape Town.

van de Ven TMFN, Fuller A, Clutton Brock T (2019a) Effects of climate change on pup growth and survival in a cooperative mammal, the meerkat. Funct Ecol 34: 194–202.

van de Ven TMFN, McKechnie AE, Cunningham SJ (2019b) The costs of keeping cool: behavioural trade-offs between foraging and thermoregulation are associated with significant mass losses in an arid-zone bird. Oecologia 191: 205–215.

van Wilgen NJ, Goodall V, Holness S, Chown SL, McGeoch MA (2016) Rising temperatures and changing rainfall patterns in South Africa’s national parks. Int J Climatol 36: 706–721.

Visser GH, Boon PE, Meijer HAJ (2000) Validation of the doubly labeled water method in Japanese Quail *Coturnix c. japonica* chicks: is there an effect of growth rate? J Comp Physiol B Biochem Syst Environ Physiol 170: 365–372.

Wada H, Kriengwatana B, Allen N, Schmidt KL, Soma KK, MacDougall-Shackleton SA (2015) Transient and permanent effects of suboptimal incubation temperatures on growth, metabolic rate, immune function and adrenocortical responses in zebra finches. J Exp Biol 218: 2847–55.

Webb DR (1987) Thermal tolerance of avian embryos: a review. Condor 89: 874–898.

Welbergen JA, Klose SM, Markus N, Eby P (2008) Climate change and the effects of temperature extremes on Australian flying-foxes. Proc R Soc B 275: 419–425.

Wiley EM, Ridley AR (2016) The effects of temperature on offspring provisioning in a cooperative breeder. Anim Behav 117: 187–195.

Williams JB, Tieleman BI (2005) Physiological adaptation in desert birds. Bioscience 55: 416–425.

