## Supporting Information Table S2 for "Dehydration risk, not ambient incubation, limits nest attendance at high temperatures"

**Running head: Dehydration limits nest attendance**

Authors: Amanda R. Bourne*[1], Amanda R. Ridley[1,2], Andrew E. McKechnie[3,4], Claire N. Spottiswoode[1,5], Susan J. Cunningham[1]

[1] FitzPatrick Institute of African Ornithology, DST-NRF Centre of Excellence, University of Cape Town, Private Bag X3, Rondebosch 7701, South Africa

[2] Centre for Evolutionary Biology, School of Biological Sciences, University of Western Australia, Crawley 6009, Australia

[3] South African Research Chair in Conservation Physiology, South African National Biodiversity Institute, Pretoria, South Africa

[4] DST-NRF Centre of Excellence at the FitzPatrick Institute, Department of Zoology and Entomology, University of Pretoria, Hatfield, South Africa

[5]Department of Zoology, University of Cambridge, Downing Street, Cambridge CB2 3EJ, UK

### Supporting Information

### Power to detect interactions

Interactions between group size and climatic effects on development would be consistent with a buffering effect of group size on survival. We therefore conducted sensitivity power analyses to identify the minimum determinable effect of two-way interactions given our sample sizes (Cohen, 1988; Greenland et al., 2016), using the package *pwr* (Champely et al., 2018). For our regression models, we used the function **pwr.f2.test(u =,v =,f2 =,sig.level =,power =)**, where u = numerator degrees of freedom, v = denominator degrees of freedom, $\alpha$ (the signficance level; probability of finding an effect that is not there) = 0.05, and power (probability of finding an effect that is there) = 0.8. The value $f^2$ is the calculated value, representing the measure of determinable effect size. We assumed a fourfold increase in required sample size to adequately detect interactions in mixed-effects models (Leon & Heo, 2009), and found that two-way interaction effects would have to be moderate to very large for us to be able to detect them in this dataset (all Cohen's $f^2 > 0.21$). We have sufficient sample size to detect a range of main effect sizes, from small to large, in all analyses (range $f^2$: 0.04 – 0.18) – see Table S1 below. Cohen (1988) suggested that $f^2$ values of ~ 0.02, ~0.15, and ~0.35 represent small, medium, and large effect sizes respectively.

**Table S1**

***Power analyses for the interactions: multiple regression power calculations***

| Analysis | *u* | *v* | *α* | *power* | $f^2$ |
|---|---|---|---|---|---|
| **Nest outcomes** | | | | | |
| Main effects | 2 | 99 | 0.05 | 0.8 | 0.081 |
| Interactions | 3 | 25 | 0.05 | 0.8 | 0.443 |
| **Nest attendance** | | | | | |
| Main effects | 2 | 46 | 0.05 | 0.8 | 0.178 |
| Interactions | 3 | 12 | 0.05 | 0.8 | 1.142 |
| **Energy expenditure and water balance** | | | | | |
| Main effects | 2 | 70 | 0.05 | 0.8 | 0.115 |
| Interactions | 3 | 17 | 0.05 | 0.8 | 0.658 |
| **Mass change** | | | | | |
| Main effects | 2 | 120 | 0.05 | 0.8 | 0.066 |
| Interactions | 3 | 30 | 0.05 | 0.8 | 0.359 |

*u = model degrees of freedom; v = sample size, α = the significance level, and power (p) = probability of finding an effect that is there; $f^2$ = measure of determinable effect size (values of ~ 0.02, ~0.15, and ~0.35 represent small, moderate, and large determinable effect sizes respectively).

### <u>Nest outcomes</u>

### Table S2

***Effects of environmental and social factors on probability of hatching***

Data from 99 nests by 23 groups over 3 breeding seasons
Random term: Nest identity
Data analysis: binomial glmer with binary response (hatch = 1, fail = 0) in *lme4*

| *Model terms* | *AICc* | *ΔAICc* | *weight* |
|---|---|---|---|
| Null model | 134.0 | 15.90 | 0.000 |
| Season | 137.7 | 19.57 | 0.000 |
| $T_{max}$ | 118.1 | 0.00 | 0.817 |
| Group size | 135.9 | 17.79 | 0.000 |
| Group size + Group size^2 | 135.8 | 17.66 | 0.000 |
| $T_{max}$ + Group size + $T_{max}$ * Group size | 121.2 | 3.00 | 0.182 |
| Season + Group size + Season * Group size | 140.4 | 22.28 | 0.000 |
| | *AICc* | *ΔAICc* | *weight* |
| Null model | 134.0 | 18.10 | 0.00 |
| *Top models:* | | | |
| $T_{max}$ | 115.9 | 0.00 | 1.00 |
| Effect size of explanatory terms after model averaging | Estimate | SE | 95% CI |
| Intercept | 10.676 | 2.669 | 5.728/16.274 |
| **$T_{max}$** | **-0.303** | **0.079** | **-0.467/-0.157** |

### Nest attendance

#### Table S3

***Effects of environmental and social factors on proportion of time that clutches were incubated***

Data from 46 observation days at 35 nests by 15 groups over 3 breeding seasons
Random term: Nest identity
Data analysis: binomial glmer with *cbind* function in *lme4*

| *Model terms* | *AICc* | *ΔAICc* | *weight* |
|---|---|---|---|
| Null model | 376.6 | 12.08 | 0.001 |
| Season | 379.4 | 14.93 | 0.000 |
| $T_{max}$ | 364.5 | 0.00 | 0.498 |
| Group size | 378.7 | 14.19 | 0.000 |
| Group size + Group size^2 | 373.8 | 9.35 | 0.005 |
| $T_{max}$ + Group size + Group size^2 | 364.6 | 0.17 | 0.457 |
| $T_{max}$ + Group size + $T_{max}$ * Group size | 369.6 | 5.14 | 0.038 |
| Season + Group size + Season * Group size | 385.0 | 20.55 | 0.000 |
| | *AICc* | *ΔAICc* | *weight* |
| Null model | 376.6 | 12.13 | 0.00 |
| *Top models:* | | | |
| $T_{max}$ | 364.47 | 0.00 | 0.520 |
| $T_{max}$ + Group size + Group size^2 | 364.64 | 0.17 | 0.480 |
| Effect size of explanatory terms after model averaging | Estimate | SE | 95% CI |
| Intercept | 6.084 | 0.855 | 4.379/7.788 |
| **$T_{max}$** | **-1.588** | **0.466** | **-2.528/-0.648** |
| Group size | 0.259 | 0.419 | -0.576/1.095 |
| Group size ^ 2 | -0.633 | 0.771 | -2.156/0.889 |

Higher temperatures were also associated with more frequent incubation recesses (Est = 0.406 ± 0.182, 95% CI: 0.039, 0.772, $z$ = 2.172; Fig. S1; Table S4), longer total durations of incubation recesses (Wilcoxon rank sum $W$ = 890, $p$ = 0.012 comparing durations between hot and cools days using 35.5˚C as the threshold; Fig. S2; Table S5), and a higher probability of observing any incubation recesses at all (Est = 1.498 ± 0.712, 95% CI: 0.061, 2.934, z = 2.043; Fig. S3; Table S6).

The babblers leave their nests unattended infrequently, averaging 2 ± 2 time a day where no group members are incubating the clutch (range: 0 – 9). After averaging the two top models (combined weight = 0.760), only $T_{max}$ significantly predicted the number of time that clutches were left unattended, with the number increasing as temperatures rose (Table S2, Figure S1). exceeding 35.5°C, identified as a critical temperature threshold in pied babblers (du Plessis et al. 2012; Wiley & Ridley 2016), the number of times that clutches were left unattended averaged 3 (± 2, range: 0 – 9), whereas on cool days (maximum temperates < 35.5°C) the number of times that clutches were left unattended averaged < 1(± 2, range: 0 – 8). We only recorded a number of times that clutches were left unattended exceeding 2 on cool days twice across three breeding seasons – both of these nests were the first nests of inexperienced pairs from groups primarily made up of siblings rather than offspring.

### Table S4

***Effects of environmental and social factors on number of times a day that clutches were left completely unattended***

Data from 46 observation days at 35 nests by 15 groups over 3 breeding seasons
Random term: Nest identity
Data analysis: poisson glmer in *lme4*

| *Model terms* | *AICc* | *ΔAICc* | *weight* |
|---|---|---|---|
| Null model | 167.9 | 4.17 | 0.057 |
| Season | 170.4 | 6.75 | 0.016 |
| $T_{max}$ | 164.6 | 0.86 | 0.300 |
| Group size | 169.6 | 5.93 | 0.024 |
| Group size + Group size^2 | 166.0 | 2.34 | 0.143 |
| $T_{max}$ + Group size + Group size^2 | 163.7 | 0.00 | 0.460 |
| | *AICc* | *ΔAICc* | *weight* |
| Null model | 167.90 | 4.20 | 0.00 |
| *Top models:* | | | |
| $T_{max}$ + Group size + Group size^2 | 163.70 | 0.00 | 0.610 |
| $T_{max}$ | 164.56 | 0.86 | 0.390 |
| Effect size of explanatory terms after model averaging | Estimate | SE | 95% CI |
| Intercept | -0.243 | 0.395 | -1.032/0.545 |
| **$T_{max}$** | **0.406** | **0.182** | **0.039/0.772** |
| Group size | -0.264 | 0.261 | -0.782/0.254 |
| Group size ^ 2 | 0.325 | 0.329 | -0.327/0.976 |

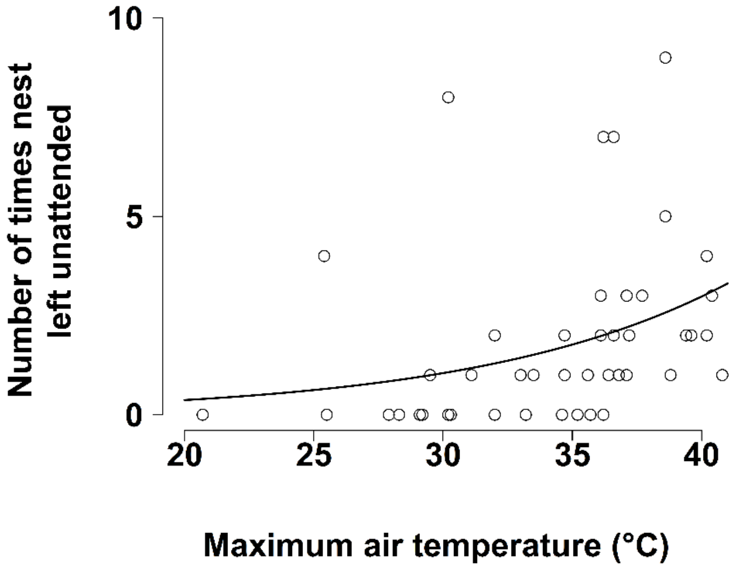


*Figure S1: Number of times that the clutch was left unattended by maximum temperature on the observation day. Data from 46 observation days at 35 nests over 3 breeding seasons.*

The duration of time periods during which clutches were left unattended was usually quite short (n = 83; mean = 22 ± 37.9 min; range: 1 – 265 min) – 80.7% of these periods were shorter than 30 min. The average duration of time periods during which clutches were left unattended was significantly longer on hot days (mean = 26.3 ± 42.6 min; range: 1 – 265 min) than on cool days (9.1 ± 10.4 min; range: 1 – 37 min; Wilcoxon rank sum $W = 890$, $p = 0.012$). Most of the time, clutches were left unattended for less than 30 min in total per day – we recorded clutch non-attendance totalling longer than 30 min per day on only 15 of 46 observation days. Thirteen of these occurred on days where maximum air temperatures exceeded 35.5°C and the other two were the nests of the inexperienced pairs from groups primarily made up of siblings rather than offspring described above. Days with no or very short total time periods for which clutches were left unattended (n = 24 with total non-attendance < 10 min) tended to be cooler (18 of 24 days had maximum temperatures < 35.5°C, mean = 33.1 ± 4.3°C; Wilcoxon rank sum $W = 104$, $p = 0.003$). This data is modelled as the inverse of nest attendance, with proportion of time that clutches were left unattended as the response and with the same results: after averaging the two top models (combined weight = 0.993), only $T_{max}$ significantly predicted the proportion of time that clutches were left unattended, with proportion time unattended increasing as temperatures rose (Table S3, Figure S2).

### Table S5

***Effects of environmental and social factors on proportion of time that clutches were left completely unattended***

Data from 46 observation days at 35 nests by 15 groups over 3 breeding seasons
Random term: Nest identity
Data analysis: binomial glmer with *cbind* function in *lme4*

| *Model terms* | *AICc* | *ΔAICc* | *weight* |
|---|---|---|---|
| Null model | 376.6 | 12.08 | 0.001 |
| Season | 379.4 | 4.93 | 0.000 |
| $T_{max}$ | 364.5 | 0.00 | 0.518 |
| Group size | 378.7 | 14.19 | 0.000 |
| Group size + Group size^2 | 373.8 | 9.35 | 0.005 |
| $T_{max}$ + Group size + Group size^2 | 364.6 | 0.17 | 0.475 |
| | *AICc* | *ΔAICc* | *weight* |
| Null model | 376.6 | 12.13 | 0.00 |
| *Top models:* | | | |
| $T_{max}$ | 364.47 | 0.00 | 0.520 |
| $T_{max}$ + Group size + Group size^2 | 364.64 | 0.17 | 0.480 |
| Effect size of explanatory terms after model averaging | Estimate | SE | 95% CI |
| Intercept | -6.084 | 0.855 | -7.788/-4.379 |
| **$T_{max}$** | **1.588** | **0.466** | **0.648/2.528** |
| Group size | -0.259 | 0.419 | -1.095/0.576 |
| Group size ^ 2 | 0.633 | 0.771 | -0.889/2.156 |

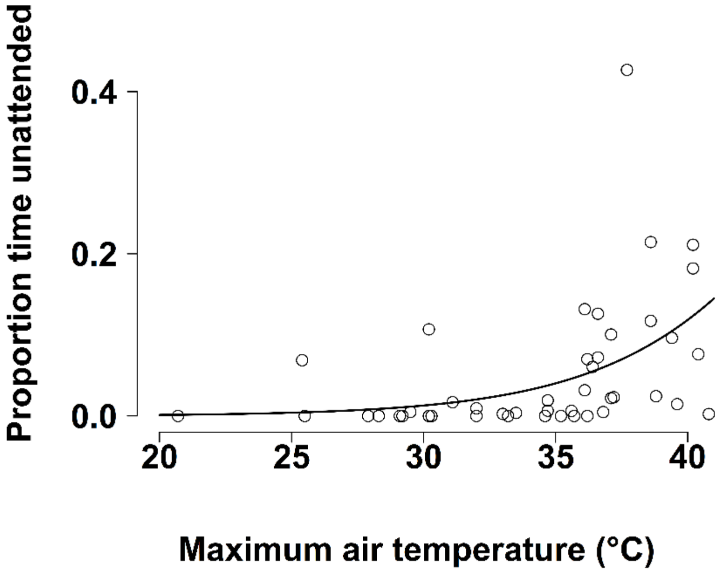


*Figure S2: Proportion of time that the clutch was left unattended by maximum temperature on the observation day. Data from 46 observation days at 35 nests over 3 breeding seasons.*

Clutches were not left unattended at all on 16 of the observation days. These days were all signficantly cooler (mean = 31.1 ± 4.3°C) than the days on which clutches were left unattended at least once (mean = 36.0 ± 3.6°C; Wilcoxon rank sum $W = 75.5$, $p = 0.001$, n = 31). After averaging the two top models (combined weight = 0.993), only $T_{max}$ significantly predicted the probability of observing that clutches were left unattended at all, with the probability of at least on period of non-attendance increasing as temperatures rose (Table S4, Figure S3). 78.6% of clutches that ultimately failed to hatch (n = 14) were left unattended at least once on our obervation days, whereas clutches that ultimately hatched (n = 21) were less likely to be left unattended - only 52.4% of were left unattended at least once on our observations days. The difference is not statistically significant ($X^2_1 = 1.473$, $p = 0.225$).

### Table S6

***Effects of environmental and social factors on the probability that clutches were left completely unattended at all***

Data from 46 observation days at 35 nests by 15 groups over 3 breeding seasons
Random term: Nest identity
Data analysis: binomial glmer in *lme4*

| *Model terms* | *AICc* | *ΔAICc* | *weight* |
|---|---|---|---|
| Null model | 62.9 | 12.10 | 0.001 |
| Season | 64.6 | 13.71 | 0.001 |
| $T_{max}$ | 51.8 | 0.92 | 0.382 |
| Group size | 64.9 | 14.01 | 0.001 |
| Group size + Group size^2 | 59.1 | 8.23 | 0.010 |
| $T_{max}$ + Group size + Group size^2 | 50.8 | 0.00 | 0.605 |
| | *AICc* | *ΔAICc* | *weight* |
| Null model | 62.90 | 12.05 | 0.00 |
| *Top models:* | | | |
| $T_{max}$ + Group size + Group size^2 | 50.85 | 0.00 | 0.610 |
| $T_{max}$ | 51.77 | 0.92 | 0.390 |
| Effect size of explanatory terms after model averaging | Estimate | SE | 95% CI |
| Intercept | 0.173 | 0.772 | -1.364/1.709 |
| **$T_{max}$** | **1.498** | **0.712** | **0.061/2.934** |
| Group size | -0.345 | 0.529 | -1.405/0.714 |
| Group size ^ 2 | 0.786 | 0.819 | -0.841/2.412 |

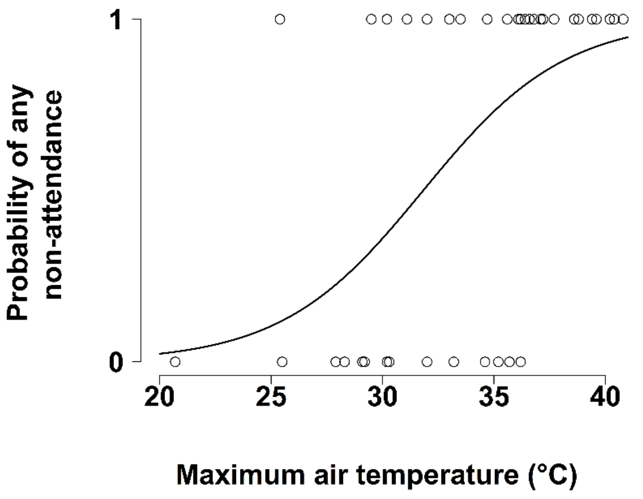


*Figure S3: Whether or not a clutch was left unattended at all by maximum temperature on the observation day. Data from 46 observation days at 35 nests over 3 breeding seasons.*

### Table S7

***Paired t.tests comparing air temperature and black bulb temperature for each hour of the day***

Data: 1,023,981 temperature records from 23 nests in *Vachellia erioloba* by 14 groups over 3 breeding seasons

| | N | Air temp mean | BB temp mean | Mann Whitney U | P value |
|---|---|---|---|---|---|
| 6 AM | 1,518 | 19.2 | 20.1 | 43932 | < 0.001 |
| 7 AM | 1,518 | 21.4 | 22.9 | 19863 | < 0.001 |
| 8 AM | 1,518 | 24.0 | 27.1 | 9600 | <0.001 |
| 9 AM | 1,518 | 26.0 | 30.3 | 6707 | <0.001 |
| 10 AM | 1,539 | 27.8 | 32.8 | 7003 | <0.001 |
| 11 AM | 1,560 | 29.2 | 33.8 | 8357 | <0.001 |
| 12 PM | 1,560 | 30.5 | 37.0 | 11664 | <0.001 |
| 1 PM | 1,560 | 31.5 | 38.2 | 11698 | <0.001 |
| 2 PM | 1,560 | 32.2 | 37.7 | 15821 | <0.001 |
| 3 PM | 1.560 | 32.6 | 36.9 | 31493 | <0.001 |
| 4 PM | 1,560 | 32.3 | 36.1 | 55564 | <0.001 |
| 5 PM | 1,554 | 32.0 | 34.0 | 98050 | <0.001 |
| 6 PM | 1,542 | 31.1 | 31.8 | 260135 | <0.001 |
| 7 PM | 1,804 | 28.9 | 29.0 | 760973 | 0.022 |

.

### Table S8

***Effects of environmental and social factors on variation in daily energy expenditure in birds from groups incubating clutches***

Data from 68 individuals from 33 nests by 15 groups over 3 breeding seasons
Random term: Bird identity
Data analysis: Gaussian lmer in lme4

| *Model terms* | *AICc* | *ΔAICc* | *weight* |
|---|---|---|---|
| Null model | 92.5 | 14.09 | 0.000 |
| Season | 93.2 | 14.75 | 0.000 |
| $T_{max}$ | 78.4 | 0.00 | 0.553 |
| Group size | 91.2 | 12.77 | 0.001 |
| Sex | 95.6 | 17.22 | 0.000 |
| Rank | 96.3 | 17.85 | 0.000 |
| $T_{max}$ + Group size | 82.4 | 3.99 | 0.075 |
| $T_{max}$ + Season | 80.0 | 1.58 | 0.251 |
| Group size + Season | 89.7 | 11.25 | 0.002 |
| $T_{max}$ + Group size + Season | 82.8 | 4.36 | 0.063 |
| $T_{max}$ + Group size + $T_{max}$ * Group size | 83.4 | 4.96 | 0.046 |
| Season + Group size + Season * Group size | 97.5 | 19.08 | 0.000 |
| $T_{max}$ + Group size + Season + $T_{max}$ * Group size | 87.1 | 8.68 | 0.007 |
| | *AICc* | *ΔAICc* | *weight* |
| Null model | 92.5 | 14.08 | 0.00 |
| *Top models:* | | | |
| $T_{max}$ | 78.42 | 0.00 | 0.69 |
| $T_{max}$ + Season | 80.00 | 1.58 | 0.31 |
| Effect size of explanatory terms | Estimate | SE | 95% CI |
| Intercept | 1.117 | 0.754 | -0.361/2.595 |
| **$T_{max}$** | **-0.223** | **0.046** | **-0.315/-0.131** |
| Season (2016-17) | 0.428 | 0.638 | -0.823/1.679 |
| Season (2017-18) | 0.550 | 0.818 | -1.054/2.155 |
| Season (2018-19) | 0.508 | 0.755 | -0.971/1.987 |

### Table S9

***Effects of environmental and social factors on variation in water balance in birds from groups incubating clutches***

Data from 69 individuals from 33 nests by 15 groups over 3 breeding seasons
Random term: Bird identity
Data analysis: Gaussian lmer in lme4

| *Model terms* | *AICc* | *ΔAICc* | *weight* |
|---|---|---|---|
| Null model | -131.0 | 0.00 | 0.945 |
| Season | -117.6 | 13.34 | 0.001 |
| $T_{max}$ | -121.6 | 9.38 | 0.009 |
| Group size | -121.4 | 9.53 | 0.008 |
| Sex | -123.2 | 7.78 | 0.019 |
| Rank | -123.0 | 7.96 | 0.018 |
| $T_{max}$ + Group size + $T_{max}$ * Group size | -102.9 | 28.09 | 0.000 |
| Season + Group size + Season * Group size | -92.0 | 38.95 | 0.000 |
| | *AICc* | *ΔAICc* | *weight* |
| *Top models:* | | | |
| Null model | -131.34 | 0.00 | 1.00 |
| Effect size of explanatory terms | Estimate | SE | 95% CI |
| Intercept | 1.025 | 0.011 | 1.004/1.045 |

### Table S10

***Effects of environmental and social factors on mass change between days in individuals from groups incubating clutches (temperature < 36.1˚C)***

Data from 72 individuals at 22 nests by 12 groups over 3 breeding seasons
Random term: Nest identity
Data analysis: Gaussian lmer in *lme4*

| *Model terms* | *AICc* | *ΔAICc* | *weight* |
|---|---|---|---|
| Null model | 317.5 | 1.12 | 0.215 |
| Season | 316.4 | 0.00 | 0.377 |
| $T_{max}$ | 320.5 | 4.13 | 0.048 |
| Group size | 320.1 | 3.71 | 0.059 |
| Sex | 319.3 | 2.91 | 0.088 |
| Rank | 319.3 | 2.92 | 0.088 |
| $T_{max}$ + Group size + $T_{max}$ * Group size | 325.5 | 9.10 | 0.004 |
| Season + Group size + Season * Group size | 318.6 | 2.27 | 0.121 |
| | *AICc* | *ΔAICc* | *weight* |
| *Top models:* | | | |
| Season | 316.36 | 0.00 | 0.640 |
| Null model | 317.48 | 1.12 | 0.36 |
| Effect size of explanatory terms | Estimate | SE | 95% CI |
| Intercept | 0.258 | 0.378 | -0.485/0.999 |
| Season (2016-17) | -0.189 | 0.509 | -1.205/0.827 |
| Season (2017-18) | 0.782 | 0.717 | -0.632/2.195 |
| Season (2018-19) | 0.507 | 0.473 | -0.425/1.439 |

### Table S11

***Effects of environmental and social factors on mass change between days in individuals from groups incubating clutches (temperature ≥ 36.1˚C)***

Data from 48 individuals at 15 nests by 10 groups over 3 breeding seasons
Random term: Nest identity
Data analysis: Gaussian lmer in *lme4*

| *Model terms* | *AICc* | *ΔAICc* | *weight* |
|---|---|---|---|
| Null model | 223.6 | 4.63 | 0.063 |
| Season | 224.5 | 5.51 | 0.040 |
| $T_{max}$ | 219.0 | 0.00 | 0.633 |
| Group size | 225.9 | 6.94 | 0.020 |
| Sex | 222.9 | 3.96 | 0.087 |
| Rank | 224.9 | 5.91 | 0.033 |
| $T_{max}$ + Group size + $T_{max}$ * Group size | 222.4 | 3.45 | 0.113 |
| Season + Group size + Season * Group size | 226.9 | 7.95 | 0.012 |
| | *AICc* | *ΔAICc* | *weight* |
| Null model | 223.6 | 5.60 | 0.00 |
| *Top model:* | | | |
| $T_{max}$ | 218.0 | 0.00 | 1.00 |
| Effect size of explanatory terms | Estimate | SE | 95% CI |
| Intercept | -0.365 | 0.315 | -1.009/-.251 |
| **$T_{max}$** | **-0.926** | **0.318** | **-1.609/-0.303** |
